# Shared genetic aetiology between cognitive functions and physical and mental health in UK Biobank (N = 112 151) and 24 GWAS consortia

**DOI:** 10.1101/031120

**Authors:** Saskia P Hagenaars, Sarah E Harris, Gail Davies, W David Hill, David CM Liewald, Stuart J Ritchie, Riccardo E Marioni, Chloe Fawns-Ritchie, Breda Cullen, Rainer Malik, METASTROKE consortium, International Consortium for Blood Pressure GWAS, SpiroMeta consortium, CHARGE consortium Pulmonary Group, CHARGE consortium Aging and Longevity Group, Bradford B Worrall, Cathie LM Sudlow, Joanna M Wardlaw, John Gallacher, Jill Pell, Andrew M McIntosh, Daniel J Smith, Catharine R Gale, Ian J Deary

## Abstract

The causes of the known associations between poorer cognitive function and many adverse neuropsychiatric outcomes, poorer physical health, and earlier death remain unknown. We used linkage disequilibrium regression and polygenic profile scoring to test for shared genetic aetiology between cognitive functions and neuropsychiatric disorders and physical health. Using information provided by many published genome-wide association study consortia, we created polygenic profile scores for 24 vascular-metabolic, neuropsychiatric, physiological-anthropometric, and cognitive traits in the participants of UK Biobank, a very large population-based sample (N = 112 151). Pleiotropy between cognitive and health traits was quantified by deriving genetic correlations using summary genome-wide association study statistics applied to the method of linkage disequilibrium regression. Substantial and significant genetic correlations were observed between cognitive test scores in the UK Biobank sample and many of the mental and physical health-related traits and disorders assessed here. In addition, highly significant associations were observed between the cognitive test scores in the UK Biobank sample and many polygenic profile scores, including coronary artery disease, stroke, Alzheimer’s disease, schizophrenia, autism, major depressive disorder, BMI, intracranial volume, infant head circumference, and childhood cognitive ability. Where disease diagnosis was available for UK Biobank participants we were able to show that these results were not confounded by those who had the relevant disease. These findings indicate that a substantial level of pleiotropy exists between cognitive abilities and many human mental and physical health disorders and traits and that it can be used to predict phenotypic variance across samples.

## Introduction

Cognitive functioning is positively associated with greater longevity and less physical and psychiatric morbidity, and negatively associated with many quantitative disease risk factors and indices.^1^ Some specific associations between cognitive functions and health appear to arise because an illness has lowered prior levels of cognitive function.^2,3^ For others, the direction of causation appears to be the reverse: there are many examples of associations between lower cognitive functions in youth, even childhood, and higher risk of later mental and physical illness and earlier death.^4-6^ In some cases, it is not clear whether illness affects cognitive functioning or vice versa, or whether both are influenced by some common factors. Many examples of these phenotypic and cognitive-illness associations are shown in Supplementary Table 1. Overall, the causes of these cognitive-health associations remain unknown and warrant further investigation. It is also well-recognised that lower educational attainment is associated with adverse health outcomes,^7^ and educational attainment has been used as a successful proxy for cognitive ability in genetic research.^8,9^ A study that included three cohorts of twins indicated that the association between higher cognitive function and increased life-span was mostly due to common genetic effects.^10^

The associations between cognitive and health and illness variables may, in part, reflect shared genetic influences. Cognitive functions show moderate heritability,^11^ and so do many physical and mental illnesses and health-associated anthropometric measures.^12^ Therefore, researchers have begun to examine pleiotropy between scores on tests of cognitive ability and health-related variables.^13^ Pleiotropy is the overlap between the genetic architecture of two or more traits, perhaps due to a variety of shared causal pathways.^14^ Originally, the possibility of pleiotropy in cognitive-health associations was tested using family- and twin-based designs.^15^ However, now data from single nucleotide polymorphism (SNP) genotyping can assess pleiotropy, opening the possibility for larger-scale, population-generalizable studies.

Multiple methods can be used to test for pleiotropy using SNP-based genetic data. Calculating genetic correlations between health measures using the summary results of previous genome-wide association studies (GWAS) has become possible using the method of linkage disequilibrium regression.^16^ In addition, the method of polygenic risk scoring^17^ also uses summary GWAS data to test whether genetic liability to a given illness or health-related anthropometric measurement is associated with phenotypes such as cognitive test scores in an second independent dataset. For example, lower cognitive functioning was associated significantly in healthy older people with higher polygenic risk for schizophrenia^18^ and stroke^19^, but not for dementia^20^. However, polygenic risk studies to date have been limited in the information they provide on this important topic: they have reported on single health outcomes, and have used relatively small cohorts with available cognitive data.

We aimed to discover whether cognitive functioning is associated with many physical and mental health and health-related anthropometric measurements in part because of their shared genetic aetiology. We curated GWAS meta-analyses for 24 health-related measures, and used them in two complementary methods to test for cognitive-health pleiotropy. First, we used linkage disequilibrium (LD) regression to derive genetic correlations between cognitive function and educational attainment traits measured in UK Biobank, and 24 health-related measures from the GWAS meta-analyses. Second, to provide a measure of the phenotypic variance that these genetic correlations account for, the summary data from GWAS meta-analyses were used to calculate polygenic risk scores in UK Biobank, which includes cognitive, educational and genome-wide SNP data on over 110 000 individuals. We calculated the associations between polygenic risk scores for the 24 health-related measures and the cognitive domains of memory, processing speed, and verbal-numerical reasoning, and educational attainment in UK Biobank participants. These new data and results provide a substantial advance in understanding the aetiology of cognitive-health associations.

## Methods

### Study design and participants

This study includes baseline data from the UK Biobank Study (http://www.ukbiobank.ac.uk).^21^ UK Biobank received ethical approval from the Research Ethics Committee (REC). The REC reference for UK Biobank is 11/NW/0382. UK Biobank is a health resource for researchers that aims to improve the prevention, diagnosis, and treatment of a range of illnesses. 502 655 community-dwelling participants aged between 37 and 73 years were recruited between 2006 and 2010 in the United Kingdom. They underwent cognitive and physical assessments, provided blood, urine, and saliva samples for future analysis, gave detailed information about their backgrounds and lifestyles, and agreed to have their health followed longitudinally. For the present study, genome-wide genotyping data were available on 112 151 individuals (58 914 female) aged 40-73 years (mean age = 56.9 years, SD = 7.9) after the quality control process (see below).

## Procedures

### Cognitive phenotypes

Three cognitive tests were used in the present study. These tests, which cover three important cognitive domains, were Reaction Time (*n* = 496 891; of whom 111 484 also had genotyping data), Memory (*n* = 498 486; 112 067 with genotyping), and Verbal-numerical Reasoning (*n* = 180 919; 36 035 with genotyping). The Reaction Time test was a computerized ‘Snap’ game, in which participants were to press a button as quickly as possible when two ‘cards’ on screen were matching. There were eight experimental trials, with a Cronbach α reliability of 0.85. In the Memory test, participants were shown a set of twelve cards (six pairs) on a computer screen for five seconds, and had to recall which were matching after the cards had been obscured. We used the number of errors in this task as the (inverse) measure of Memory ability. The Verbal-numerical Reasoning task involved a series of thirteen items assessing verbal and arithmetical deduction (Cronbach α reliability = 0.62). 492 513 participants also reported whether or not they had a college or university degree (henceforth referred to as ‘Educational Attainment’), 111 114 of whom had genotyping data. Full details on the content and administration of each test (and the Educational Attainment question) are provided in the Supplementary Materials. Supplementary Figure 1 shows that the broad age distribution of the cognitive tests did not differ between the full and genotyped samples.

### Genotyping and quality control

152 729 UK Biobank blood samples were genotyped using either the UK BiLEVE array (N = 49 979)^22^ or the UK Biobank axiom array (N = 102 750). Details of the array design, genotyping, quality control and imputation are available in a publication^22^ and in the Supplementary Materials. Quality control was performed by Affymetrix, the Wellcome Trust Centre for Human Genetics, and by the present authors; this included removal of participants based on missingness, relatedness, gender mismatch, non-British ancestry, and other criteria, and is described in the Supplementary Materials.

### Genome-wide association analyses (GWAS) in the UK Biobank sample

GWAS analyses were performed on the three UK Biobank cognitive test scores and on the Educational Attainment data in order to use the summary results for linkage disequilibrium regression. Details of the GWAS procedures are provided in the Supplementary Materials.

### Curation of summary results from GWAS consortia on health-related variables

In order to conduct LD score regression and polygenic profile score analyses between the UK Biobank cognitive data and the genetic predisposition to a large number of health-related variables, selected because of previous associations with cognitive functions (Supplementary Table 1), we gathered 24 sets of summary results from international GWAS consortia. Details of the health-related variables, the consortia’s websites, key references, and number of subjects included in each consortia’s genome-wide association study are given in Supplementary Table 2.

## Statistical Analysis

### Computing genetic associations between health and cognitive variables

We used two methods to compute genetic associations between health variables from 24 GWAS consortia, and cognitive and Educational Attainment variables measured in UK Biobank: LD score regression and polygenic profile/risk scoring. Each provides a different metric to infer the existence of pleiotropy between pairs of traits. LD score regression was used to derive genetic correlations to determine the degree to which the polygenic architecture of a trait overlaps with that of another. Next, the polygenic risk score method was used to test the extent to which these genetic correlations are predictive of phenotypic variance across samples. Both LD score regression and polygenic risk scores are dependent on the traits analysed being highly polygenic in nature, i.e. where a large number of variants of small effect contribute toward phenotypic variation.^16,17^ This was tested for each trait prior to running the LD score regression and polygenic risk analyses.

### LD score regression

In order to quantify the level of pleiotropy between the traits assessed here, LD score regression was used.^13,16^ LD score regression is a class of techniques that exploits the correlational structure of the single nucleotide polymorphisms (SNPs) found across the genome. In Supplementary Materials we provide more details of LD score regression. Here, we use LD score regression to derive genetic correlations between health-related and cognitive traits using 24 large GWAS consortia data sets that enable pleiotropy of their health-related traits to be quantified with the cognitive traits in UK Biobank. We followed the data processing pipeline devised by Bulik-Sullivan et al.^16^, described in more detail in the Supplementary Materials. In order to ensure that the genetic correlation for the Alzheimer’s disease phenotype was not driven by a single locus or biased the fit of the regression model, a 500kb region centred on the *APOE* locus was removed and this phenotype was re-run. This additional model is referred to in the Tables and Figures below as ‘Alzheimer’s disease (500kb)’.

### Polygenic profiling

The UK Biobank genotyping data required recoding from numeric (1, 2) allele coding to standard ACGT format prior to being used in polygenic profile scoring analyses. This was achieved using a bespoke programme developed by one of the present authors (DCML), details of which are provided in the Supplementary Materials.

Polygenic profiles were created for 24 health-related phenotypes (see Table 3, and Supplementary Table 1) in all genotyped participants using PRSice.^23^ PRSice calculates the sum of alleles associated with the phenotype of interest across many genetic loci, weighted by their effect sizes estimated from a GWAS of that phenotype in an independent sample. Prior to creating the scores, SNPs with a minor allele frequency < 0.01 were removed and clumping was used to obtain SNPs in linkage equilibrium with an r^2^ < 0.25 within a 200bp window. Multiple scores were then created for each phenotype containing SNPs selected according to the significance of their association with the phenotype. The GWAS summary data for each of the 24 health-related phenotypes were used to create five polygenic profiles in the UK Biobank participants, at thresholds of p < 0.01, p < 0.05, p < 0.1, p < 0.5 and all SNPs.

Correlation coefficients were calculated between each of the UK Biobank cognitive phenotypes. The associations between the polygenic profiles and the target phenotype were examined in regression models (linear regression for the continuous cognitive traits, and logistic regression for the binary education variable), adjusting for age at measurement, sex, genotyping batch and array, assessment centre, and the first ten genetic principal components to adjust for population stratification. We corrected for multiple testing across all polygenic profile scores at all significance thresholds for associations with all cognitive phenotypes (470 tests) using the False Discovery Rate (FDR) method.^24^ We conducted sensitivity analyses as follows. Where the original findings were FDR significant, UK Biobank participants with cardiovascular disease (N=5300) were then removed from analyses of coronary artery disease, those with diabetes (N=5800) were removed from type 2 diabetes analyses, and those with hypertension (N=26 912) were removed from systolic blood pressure analyses. See Supplementary Materials for further details of these sensitivity analysis. Four multivariate regression models were then performed, including all 24 polygenic profile scores and the covariates described above.

## Results

### Phenotypic and genetic associations among the UK Biobank’s cognitive traits

In addition to descriptive statistics, Table 1 shows phenotypic correlations and genetic correlations (calculated using LD score regression) among the cognitive traits in UK Biobank. There were modest correlations between the three cognitive test scores; those who did well on one test tended to do well on the other two. Verbal-numerical Reasoning showed the highest phenotypic correlation with Educational Attainment (r = 0.30). The strongest genetic correlation was also between Verbal-numerical Reasoning and Educational Attainment (r_g_ = 0.79). All but one of the other genetic correlations were statistically significant: there was a non-significant genetic correlation between Educational Attainment and Reaction Time. These results show that different cognitive traits and Educational Attainment correlate, in part, due to overlapping genetic architecture.

**Table 1.** Descriptive statistics and phenotypic (below diagonal) and genetic (above diagonal and shaded) correlations for the UK Biobank cognitive and educational variables. Genetic correlations are based on the results of genome-wide association studies of the UK Biobank variables. P-values for the correlations are shown in parentheses. For the phenotypic variables, Pearson correlations used for continuous-continuous correlations and point-biserial correlations for continuous-categorical correlations. All variables are coded such that higher scores indicate better performance. ^*^, p-value < 0.0001; ^†^ pvalue < 0.05

| Variable | Mean (SD) | Phenotypic/genetic correlation |  |  |  |
| --- | --- | --- | --- | --- | --- |
|  |  | Reaction Time | Memory | Verbal-numerical Reasoning | Educational Attainment |
| Reaction Time (ms) | 555.08 (112.69) | - | 0.179* | 0.206* | 0.066 |
| Memory (errors) | 4.06 (3.23) | 0.140* | - | 0.436* | 0.126† |
| Verbal-numerical Reasoning<br>(max. score 13) | 6.16 (2.10) | 0.179* | 0.186* | - | 0.792* |
| Educational Attainment | 30.5% with degree | 0.081* | 0.046* | 0.301* | - |

### Cognitive-health pleiotropy: overview

To test for pleiotropy between cognitive and health traits, we present the LD score regression (Table 2, Figure 1) and polygenic risk score (Table 3, Supplementary Tables 4a-d) results for the four UK Biobank cognitive function and Educational Attainment traits, and the 24 health-related traits from GWAS consortia. Results are FDR-corrected for multiple comparisons.

**Table 2.**
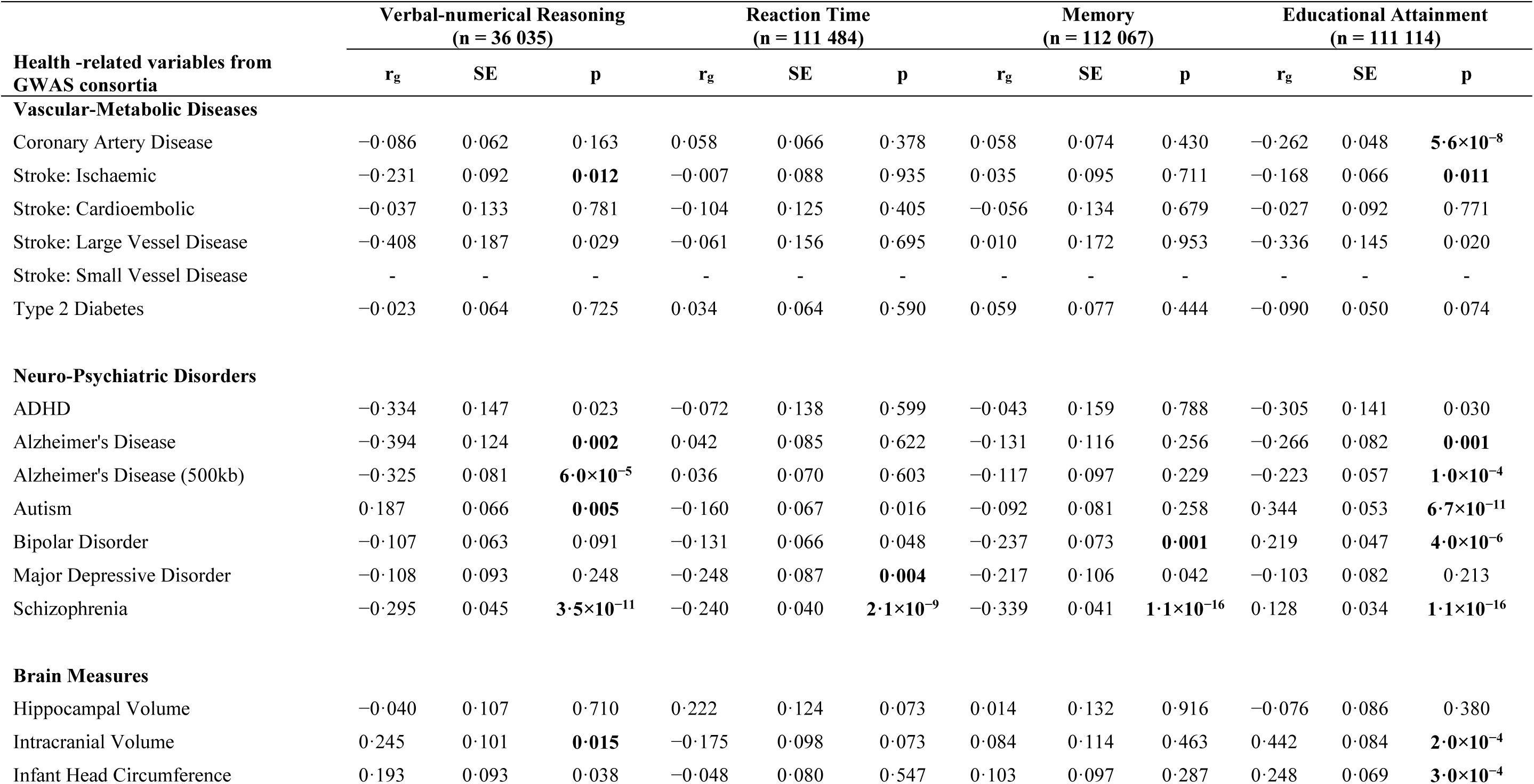
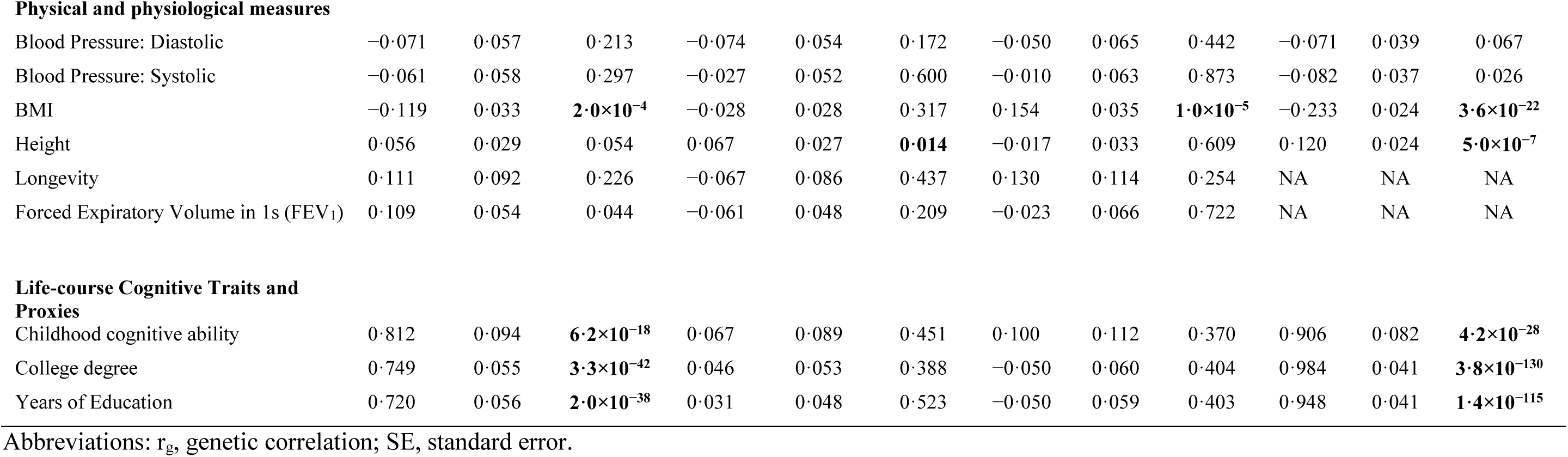
Genetic correlations between the cognitive and education phenotypes documented in the UK Biobank data set and the health-related variables collected from GWAS consortia. Statistically significant p-values (after False Discovery Rate correction; threshold: p < 0.016) are shown in bold. There was no evidence for a sufficient polygenic signal in the small vessel disease data set and so no genetic correlation could be derived as shown in supplementary Table 3.

| Health -related variables from GWAS consortia | Verbal-numerical Reasoning<br>(n = 36 035) |  |  | Reaction Time<br>(n = 111 484) |  |  | Memory<br>(n = 112 067) |  |  | Educational Attainment<br>(n = 111 114) |  |  |
| --- | --- | --- | --- | --- | --- | --- | --- | --- | --- | --- | --- | --- |
| | $r_g$ | SE | p | $r_g$ | SE | p | $r_g$ | SE | p | $r_g$ | SE | p |
| **Vascular-Metabolic Diseases** |  |  |  |  |  |  |  |  |  |  |  |  |
| Coronary Artery Disease | -0.086 | 0.062 | 0.163 | 0.058 | 0.066 | 0.378 | 0.058 | 0.074 | 0.430 | -0.262 | 0.048 | **$5.6 \times 10^{-8}$** |
| Stroke: Ischaemic | -0.231 | 0.092 | **0.012** | -0.007 | 0.088 | 0.935 | 0.035 | 0.095 | 0.711 | -0.168 | 0.066 | **0.011** |
| Stroke: Cardioembolic | -0.037 | 0.133 | 0.781 | -0.104 | 0.125 | 0.405 | -0.056 | 0.134 | 0.679 | -0.027 | 0.092 | 0.771 |
| Stroke: Large Vessel Disease | -0.408 | 0.187 | 0.029 | -0.061 | 0.156 | 0.695 | 0.010 | 0.172 | 0.953 | -0.336 | 0.145 | 0.020 |
| Stroke: Small Vessel Disease | - | - | - | - | - | - | - | - | - | - | - | - |
| Type 2 Diabetes | -0.023 | 0.064 | 0.725 | 0.034 | 0.064 | 0.590 | 0.059 | 0.077 | 0.444 | -0.090 | 0.050 | 0.074 |
| **Neuro-Psychiatric Disorders** |  |  |  |  |  |  |  |  |  |  |  |  |
| ADHD | -0.334 | 0.147 | 0.023 | -0.072 | 0.138 | 0.599 | -0.043 | 0.159 | 0.788 | -0.305 | 0.141 | 0.030 |
| Alzheimer's Disease | -0.394 | 0.124 | **0.002** | 0.042 | 0.085 | 0.622 | -0.131 | 0.116 | 0.256 | -0.266 | 0.082 | **0.001** |
| Alzheimer's Disease (500kb) | -0.325 | 0.081 | **$6.0 \times 10^{-5}$** | 0.036 | 0.070 | 0.603 | -0.117 | 0.097 | 0.229 | -0.223 | 0.057 | **$1.0 \times 10^{-4}$** |
| Autism | 0.187 | 0.066 | **0.005** | -0.160 | 0.067 | 0.016 | -0.092 | 0.081 | 0.258 | 0.344 | 0.053 | **$6.7 \times 10^{-11}$** |
| Bipolar Disorder | -0.107 | 0.063 | 0.091 | -0.131 | 0.066 | 0.048 | -0.237 | 0.073 | **0.001** | 0.219 | 0.047 | **$4.0 \times 10^{-6}$** |
| Major Depressive Disorder | -0.108 | 0.093 | 0.248 | -0.248 | 0.087 | **0.004** | -0.217 | 0.106 | 0.042 | -0.103 | 0.082 | 0.213 |
| Schizophrenia | -0.295 | 0.045 | **$3.5 \times 10^{-11}$** | -0.240 | 0.040 | **$2.1 \times 10^{-9}$** | -0.339 | 0.041 | **$1.1 \times 10^{-16}$** | 0.128 | 0.034 | **$1.1 \times 10^{-16}$** |
| **Brain Measures** |  |  |  |  |  |  |  |  |  |  |  |  |
| Hippocampal Volume | -0.040 | 0.107 | 0.710 | 0.222 | 0.124 | 0.073 | 0.014 | 0.132 | 0.916 | -0.076 | 0.086 | 0.380 |
| Intracranial Volume | 0.245 | 0.101 | **0.015** | -0.175 | 0.098 | 0.073 | 0.084 | 0.114 | 0.463 | 0.442 | 0.084 | **$2.0 \times 10^{-4}$** |
| Infant Head Circumference | 0.193 | 0.093 | 0.038 | -0.048 | 0.080 | 0.547 | 0.103 | 0.097 | 0.287 | 0.248 | 0.069 | **$3.0 \times 10^{-4}$** |
| **Physical and physiological measures** |  |  |  |  |  |  |  |  |  |  |  |  |
| Blood Pressure: Diastolic | -0.071 | 0.057 | 0.213 | -0.074 | 0.054 | 0.172 | -0.050 | 0.065 | 0.442 | -0.071 | 0.039 | 0.067 |
| Blood Pressure: Systolic | -0.061 | 0.058 | 0.297 | -0.027 | 0.052 | 0.600 | -0.010 | 0.063 | 0.873 | -0.082 | 0.037 | 0.026 |
| BMI | -0.119 | 0.033 | $2.0 \times 10^{-4}$ | -0.028 | 0.028 | 0.317 | 0.154 | 0.035 | $1.0 \times 10^{-5}$ | -0.233 | 0.024 | $3.6 \times 10^{-22}$ |
| Height | 0.056 | 0.029 | 0.054 | 0.067 | 0.027 | **0.014** | -0.017 | 0.033 | 0.609 | 0.120 | 0.024 | $5.0 \times 10^{-7}$ |
| Longevity | 0.111 | 0.092 | 0.226 | -0.067 | 0.086 | 0.437 | 0.130 | 0.114 | 0.254 | NA | NA | NA |
| Forced Expiratory Volume in 1s (FEV<sub>1</sub>) | 0.109 | 0.054 | 0.044 | -0.061 | 0.048 | 0.209 | -0.023 | 0.066 | 0.722 | NA | NA | NA |
| **Life-course Cognitive Traits and Proxies** |  |  |  |  |  |  |  |  |  |  |  |  |
| Childhood cognitive ability | 0.812 | 0.094 | $6.2 \times 10^{-18}$ | 0.067 | 0.089 | 0.451 | 0.100 | 0.112 | 0.370 | 0.906 | 0.082 | $4.2 \times 10^{-28}$ |
| College degree | 0.749 | 0.055 | $3.3 \times 10^{-42}$ | 0.046 | 0.053 | 0.388 | -0.050 | 0.060 | 0.404 | 0.984 | 0.041 | $3.8 \times 10^{-130}$ |
| Years of Education | 0.720 | 0.056 | $2.0 \times 10^{-38}$ | 0.031 | 0.048 | 0.523 | -0.050 | 0.059 | 0.403 | 0.948 | 0.041 | $1.4 \times 10^{-115}$ |

Abbreviations: $r_g$ , genetic correlation; SE, standard error.

**Figure 1.**
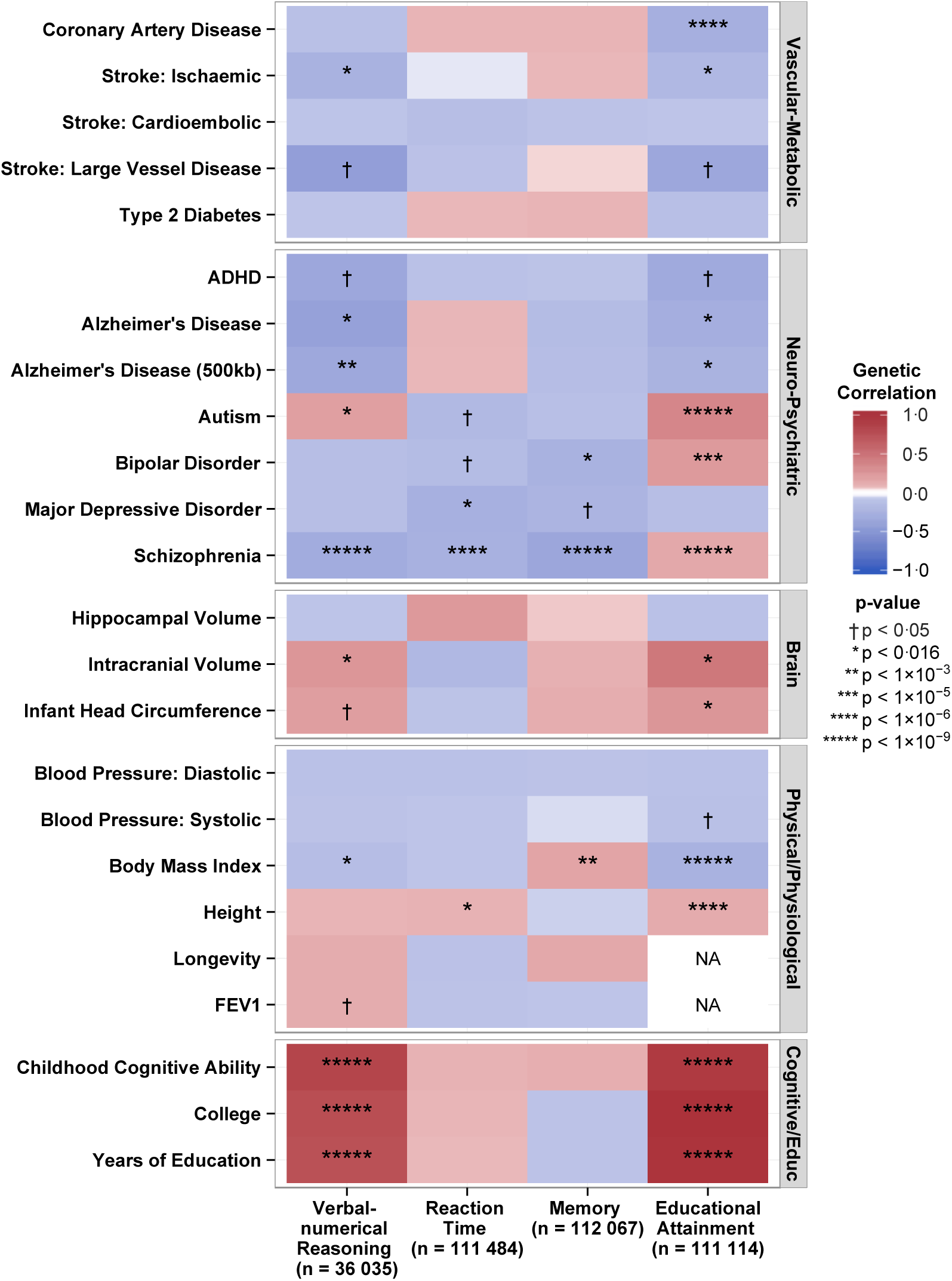
Heat map of genetic correlations calculated using LD regression between cognitive phenotypes in UK Biobank and health-related variables from GWAS consortia. Hues and colours depict, respectively, the strength and direction of the genetic correlation between the cognitive phenotypes in UK Biobank and the health-related variables. Red and blue indicate positive and negative correlations, respectively. Correlations with the darker shade associated with a stronger association. Based on results in Table 2

**Table 3.** Associations between polygenic profiles of health related traits created from GWAS consortia summary data, and UK Biobank cognitive and education phenotypes controlling for age, sex, assessment centre, genotyping batch and array, and ten principal components for population structure. FDR-corrected statistically significant values (P<0.0188) are shown in bold. Cognitive and education phenotypes are scored such that higher scores indicate better performance. The associations between the polygenic profile with the largest effect size (thresh) and each cognitive and education phenotype are presented. Thresh; the p-value threshold with the largest effect size.

| Health -related variables<br>from GWAS consortia | Verbal-Numerical Reasoning<br>(n = 36 035) |  |  | Reaction Time<br>(n = 111 484) |  |  | Memory<br>(n = 112 067) |  |  | Educational Attainment<br>(n = 111 114) |  |  |
| --- | --- | --- | --- | --- | --- | --- | --- | --- | --- | --- | --- | --- |
| | thresh | $\beta$ | p | thresh | $\beta$ | p | thresh | B | p | thresh | $\beta$ | p |
| **Vascular-Metabolic Diseases** |  |  |  |  |  |  |  |  |  |  |  |  |
| Coronary Artery Disease | 0.5 | -0.019 | **0.0002<sup>a</sup>** | 0.1 | 0.001 | 0.0875 | 0.1 | -0.004 | 0.2265 | 1 | -0.047 | **7.92×10<sup>-13b</sup>** |
| Stroke: Ischaemic | 0.05 | -0.014 | **0.0068** | 0.01 | -0.001 | 0.0791 | 0.01 | -0.004 | 0.1759 | 0.5 | -0.020 | **0.0026** |
| Stroke: Cardioembolic | 1 | -0.009 | 0.0937 | 0.01 | 0.001 | 0.0817 | 0.01 | 0.005 | 0.0638 | 0.5 | -0.003 | 0.6931 |
| Stroke: Large Vessel Disease | 0.01 | -0.013 | **0.0155** | 0.05 | 2.26×10<sup>-4</sup> | 0.6714 | 0.05 | -0.008 | **0.0056** | 0.1 | -0.031 | **5.31×10<sup>-6</sup>** |
| Stroke: Small Vessel Disease | 0.05 | -0.012 | 0.0250 | 1 | -0.001 | 0.3188 | 1 | -0.004 | 0.1388 | 0.1 | -0.034 | **2.96×10<sup>-7</sup>** |
| Type 2 Diabetes | 1 | -0.006 | 0.2490 | 0.1 | -4.50×10<sup>-4</sup> | 0.3982 | 0.1 | 0.003 | 0.2614 | 0.5 | -0.025 | **0.0002<sup>c</sup>** |
| **Neuro-Psychiatric Disorders** |  |  |  |  |  |  |  |  |  |  |  |  |
| ADHD | 1 | -0.008 | 0.1054 | 1 | -0.001 | 0.2485 | 1 | 0.001 | 0.6520 | 1 | -0.027 | **4.68×10<sup>-5</sup>** |
| Alzheimer's Disease | 0.05 | -0.023 | **1.27×10<sup>-5</sup>** | 0.5 | -0.001 | 0.0516 | 0.5 | -0.011 | **1.22×10<sup>-4</sup>** | 1 | -0.046 | **2.33×10<sup>-12</sup>** |
| Autism | 1 | 0.023 | **1.43×10<sup>-5</sup>** | 0.1 | 0.001 | 0.1176 | 0.1 | 0.002 | 0.4385 | 0.5 | 0.068 | **5.00×10<sup>-25</sup>** |
| Bipolar Disorder | 0.01 | -0.006 | 0.2695 | 0.01 | -0.002 | **0.0007** | 0.01 | -0.017 | **3.21×10<sup>-8</sup>** | 0.5 | 0.057 | **1.35×10<sup>-17</sup>** |
| Major Depressive Disorder | 1 | -0.020 | **0.0002** | 0.1 | -0.002 | **0.0005** | 0.1 | -0.014 | **1.91×10<sup>-6</sup>** | 1 | -0.009 | 0.1905 |
| Schizophrenia | 0.05 | -0.062 | **7.73×10<sup>-32</sup>** | 0.1 | -0.006 | **1.88×10<sup>-25</sup>** | 0.1 | -0.039 | **4.49×10<sup>-39</sup>** | 0.05 | 0.025 | **0.0002** |
| **Brain Measures** |  |  |  |  |  |  |  |  |  |  |  |  |
| Hippocampal Volume | 0.01 | 0.005 | 0.3552 | 0.5 | 0.001 | 0.1144 | 0.5 | -0.005 | 0.0848 | 0.5 | 0.007 | 0.2887 |
| Intracranial Volume | 0.5 | 0.016 | **0.0021** | 0.01 | 0.001 | 0.3306 | 0.01 | 0.005 | 0.0778 | 1 | 0.043 | **2.91×10<sup>-10</sup>** |
| Infant Head Circumference | 0.5 | 0.024 | **8.53×10<sup>-6</sup>** | 0.01 | -0.001 | 0.0468 | 0.01 | 0.005 | 0.0782 | 0.1 | 0.050 | **4.01×10<sup>-14</sup>** |
| **Physical and physiological measures** |  |  |  |  |  |  |  |  |  |  |  |  |
| Blood Pressure: Diastolic | 0.05 | 0.004 | 0.5044 | 0.01 | 0.001 | 0.2325 | 0.01 | -0.002 | 0.5342 | 0.1 | -0.022 | **0.0011** |
| Blood Pressure: Systolic | 0.1 | 0.013 | **0.0116** | 0.5 | $-4.97 \times 10^{-4}$ | 0.3460 | 0.5 | -0.003 | 0.3149 | 0.1 | -0.035 | **$7.61 \times 10^{-8d}$** |
| BMI | 0.5 | -0.027 | **$3.34 \times 10^{-7}$** | 0.01 | $3.96 \times 10^{-4}$ | 0.4507 | 0.01 | 0.016 | **$7.31 \times 10^{-8}$** | 0.5 | -0.093 | **$6.40 \times 10^{-45}$** |
| Height | 1 | 0.021 | **$9.23 \times 10^{-5}$** | 0.5 | $4.46 \times 10^{-4}$ | 0.4135 | 0.5 | 0.003 | 0.3579 | 1 | 0.070 | **$2.95 \times 10^{-24}$** |
| Longevity | 0.01 | 0.004 | 0.4091 | 0.05 | $3.23 \times 10^{-4}$ | 0.5426 | 0.05 | 0.006 | 0.0484 | NA | NA | NA |
| Forced Expiratory Volume in 1s (FEV<sub>1</sub>) | 0.5 | 0.012 | 0.0220 | 0.05 | -0.001 | 0.1302 | 0.05 | -0.005 | 0.0840 | NA | NA | NA |
| **Life-course Cognitive Traits and Proxies** |  |  |  |  |  |  |  |  |  |  |  |  |
| Childhood cognitive ability | 1 | 0.079 | **$3.49 \times 10^{-51}$** | 0.5 | 0.002 | **0.0002** | 0.5 | 0.014 | **$1.65 \times 10^{-6}$** | 1 | 0.122 | **$3.92 \times 10^{-75}$** |
| College degree | 0.5 | 0.102 | **$3.19 \times 10^{-83}$** | 0.5 | 0.001 | **0.0133** | 0.5 | -0.002 | 0.4688 | 0.5 | 0.280 | **0** |
| Years of Education | 1 | 0.106 | **$2.96 \times 10^{-89}$** | 0.1 | 0.001 | **0.0156** | 0.1 | -0.004 | 0.1563 | 1 | 0.210 | **0** |

<sup>a</sup> Excluding individuals with cardiovascular disease $\beta=-0.018$ , $p=0.0007$ ; <sup>b</sup> excluding individuals with cardiovascular disease $\beta=-0.044$ , $p=4.66 \times 10^{-11}$ ; <sup>c</sup> excluding individuals with diabetes $\beta=-$
$0.020$ , $p=0.0032$ ; <sup>d</sup> excluding individuals with hypertension $\beta=-0.028$ , $p=0.00016$

In overview, using LD score regression results, Educational Attainment showed significant genetic correlations with 14 of the 22 health-related traits, Verbal-numerical Reasoning had significant genetic correlations with ten of the 24, and Memory and Reaction Time both correlated significantly with three of the 24 (Table 2, Figure 1).

In the polygenic profile score results, using the best SNP threshold of the five that were created, summary GWAS data from 19 of 22 consortia predicted significant phenotypic variation in Educational Attainment in the UK Biobank sample (Table 3). For Verbal-numerical Reasoning, results from 15 of the 24 consortia predicted significant phenotypic variation. The numbers for Memory and Reaction Time were seven and six, respectively. The numbers of SNPs included in each polygenic threshold score for each of the 24 health-related traits are shown in Supplementary Table 3. The fuller results relating all five of the SNP thresholds for all 24 health-associated variables with the UK Biobank cognitive phenotypes are shown in Supplementary Tables 4a-d.

### Cognitive-health pleiotropy: brain-cognitive traits

Cognitive and brain traits provided a first check prior to moving to more mainstream health measures whose phenotypes appear more distant from cognitive function. In LD score regression analyses we expected significant genetic correlations between GWAS consortia results from cognitive- and brain-related traits and the UK Biobank GWAS results for cognitive variables. In polygenic risk score analyses we expected the polygenic scores based on the health-related GWAS consortia’s results to predict phenotypic variation in UK Biobank’s cognitive traits.

Using LD score regression, genetic correlations of greater than 0.9 were found between UK Biobank’s Education Attainment variable and GWAS consortia results from childhood cognitive ability, college degree attainment, and years of education (Table 2, Figure 1). The genetic correlation between UK Biobank’s Educational Attainment variable and ENIGMA’s intracranial volume was 0.44 and with Early Growth Genetics Consortium’s (EGG) infant head circumference was 0.25. Verbal-numerical reasoning in UK Biobank had a genetic correlation of 0.81 with childhood cognitive ability, of 0.75 with attaining a college degree from the Social Science Genetic Association consortium (SSGAC), 0.72 with years of education (SSGAC), and 0.24 with intracranial volume. These results demonstrate substantial shared genetic aetiology between brain size, cognitive ability, and educational attainment. Reaction time and memory did not have significant genetic correlations with the brain, cognitive or educational variables.

In summarising polygenic profile analyses’ results in the text we shall use standardised betas (β). Polygenic profiles for higher childhood cognitive ability and a higher level of educational attainment (SSGAC) were significantly associated with increased likelihood of a college degree (β between 0.12 and 0.28), higher scores on verbal numerical reasoning (β between 0.08 and 0.11), and faster reaction time (β about 0.001). Only childhood cognitive ability was significantly associated with memory (β = 0.01). Polygenic profiles for greater intracranial volume and greater infant head circumference were significantly associated with increased likelihood of a college degree (β = 0.04 and 0.05, respectively), and higher scores on verbal numerical reasoning (both β = 0.02).

### Cognitive-health pleiotropy: neuro-psychiatric disorders

Using LD score regression, there were negative genetic correlations between Alzheimer’s disease and UK Biobank’s Educational Attainment variable and Verbal-numerical Reasoning (r_g_ between −0.27 and −0.39) (Table 2, Figure 1). Autism had positive genetic correlations with the same two Biobank variables: 0.34 for Educational Attainment and 0.19 for Verbal-numerical Reasoning. Bipolar disorder and schizophrenia showed a pattern of having a positive genetic correlation with Educational Attainment (0.22 and 0.13, respectively) and a negative genetic correlation with verbal-numerical reasoning (−0.11 and −0.30, respectively). Schizophrenia was genetically associated with slower reaction time (−0.24) and poorer memory (−0.34). Bipolar disorder was also genetically associated with poorer memory (−0.24). There were no significant genetic correlations with major depressive disorder and any UK Biobank cognitive trait.

Polygenic profile scoring replicated the directions of association found with LD score regression-estimated genetic correlations (Table 3). Higher polygenic risk for Alzheimer’s disease was associated with lower Educational Attainment and a lower score on verbal-numerical reasoning (β between −0.01 and −0.05). Higher polygenic risk for ADHD was associated with lower Educational Attainment (β = 0.03). Higher polygenic risk for autism was associated with higher Educational Attainment and better Verbal-numerical Reasoning (β = 0.07 and 0.02, respectively). Higher polygenic risk for bipolar disorder and schizophrenia had a positive association with Educational Attainment (β = 0.06 and 0.03, respectively), and the genetic risk for schizophrenia and major depressive disorder had a negative association with Verbal-numerical Reasoning (β = −0.06 and −0.02, respectively).

### Cognitive-health pleiotropy: vascular-metabolic diseases

There were significant negative genetic correlations between UK Biobank’s Educational Attainment variable and coronary artery disease (−0.26), ischaemic stroke (−0.17), and large vessel disease stroke (−0.34) (Table 2, Figure 1). There was a significant negative genetic correlation between ischaemic stroke and Verbal-numerical Reasoning (−0.23).

Greater polygenic risk for coronary artery disease, type 2 diabetes, and ischaemic, and large and small vessel disease stroke were all associated with lower Educational Attainment (β between −0.02 and −0.05). Greater polygenic risk for coronary artery disease, and ischaemic and large vessel disease stroke were associated with lower Verbal-numerical Reasoning scores (β between −0.01 and −0.02). Greater polygenic risk for large vessel disease stroke was associated with more errors in the Memory task (β = −0.01). There was little change to the results for coronary artery disease when individuals diagnosed with cardiovascular disease were removed from the analyses, and little change for type 2 diabetes results when individuals with diabetes were removed (Table 3).

### Cognitive-health pleiotropy: physical, physiological and anthropometric measures

There were significant genetic correlations between UK Biobank’s Educational Attainment variable and BMI (−0.23), and height (0.12), both obtained from the GIANT consortium. There was a significant negative genetic correlation between BMI and Verbal-numerical Reasoning (−0.12) and a significant positive genetic correlation with Memory (0.15).

Greater polygenic risk for diastolic and systolic blood pressure (obtained from ICBP), and BMI were all associated with lower Educational Attainment (β of −0.02, −0.04, and −0.09, respectively). A polygenic profile for greater height was associated with higher Educational Attainment (β = 0.07). Higher Verbal-numerical Reasoning scores were associated with lower polygenic risk for BMI (β = −0.03), and a polygenic profile for greater height (β = 0.02) and higher systolic blood pressure (β = 0.01). Results for systolic blood pressure changed little when individuals with hypertension were removed from the analysis (Table 3).

### Multivariate models predicting cognitive variance using many polygenic profile scores

We next ran four multivariate regression models that included all 24 polygenic profile scores alongside the same covariates as were described above. This tested whether there was redundancy amongst the polygenic profile scores, and the extent to which including them all together in a multivariate model would improve the prediction of the cognitive phenotype. We compared the *R*^2^ value of models including all the profile scores to models including only the covariates. The polygenic profile scores alone accounted for 3.33% of the variance in Educational Attainment of a college or university degree, 2.26% of the variance in Verbal-numerical Reasoning scores, 0.12% in Reaction Time, and 0.16% in Memory scores. See Supplementary Table 5 for full results.

## Discussion

The present study has combined the power of UK Biobank’s very large genotyped and cognitively-tested sample with the summary results of 24 large international GWAS consortia of physical and mental disorders and health-related traits. Two methods—LD score regression and polygenic profile scoring based on previous GWAS findings—discovered extensive cognitive-health pleiotropy and showed that it can be used to predict phenotypic variance between GWAS data sets. Our results provide comprehensive new findings on the overlaps between phenotypic cognitive ability levels, genetic bases for health-related characteristics such as height and blood pressure, and liabilities to physical and psychiatric disorders even in mostly healthy, non-diagnosed individuals. They make important steps toward understanding the specific patterns of overlap between biological influences on health and their consequences for key cognitive abilities. For example, some of the association between educational attainment—often used as a social background indicator—and health appears to have a genetic aetiology. These results should stimulate further research that will be informative about the specific genetic mechanisms of the associations found here, which likely involves both protective and detrimental effects of different genetic variants.

Findings for polygenic risk for coronary artery disease were not confounded by individuals with a diagnosis of cardiovascular disease, findings for type 2 diabetes were not confounded by individuals with a diagnosis of diabetes, and findings for systolic blood pressure were not confounded by individuals with a diagnosis of hypertension. These results indicate that even in healthy individuals, being at high polygenic risk for coronary artery disease, type 2 diabetes or high blood pressure is associated with lower cognitive function and lower educational attainment.

Using LD score regression, we quantified for the genetic correlations from molecular genetic evidence between tests of cognitive ability and a wealth of health and anthropometric traits in over 100 000 individuals. As shown in Table 2, Verbal-numerical Reasoning and Educational Attainment showed a greater degree of pleiotropy than Reaction Time and Memory, with many of the health and anthropometric variables studied here. Novel genetic correlations were quantified between cognitive function, using Verbal-numerical Reasoning, and schizophrenia, Alzheimer’s disease, and ischaemic stroke. It has not escaped our notice that there are multiple possible interpretations of these genetic correlations. Not only might particular genes contribute both to cognitive and health-related traits, but genetic variants relating to health conditions could have indirect effects on cognitive ability (e.g., via medications used to treat disorders), and vice versa (e.g., via cognitively-associated lifestyle choices). See Solovieff et al.^14^ for discussion of these issues of causality and pleiotropy.

Perhaps counter-intuitively, our results indicated that the genetic variants associated with obtaining a college degree were also related to higher genetic risk of schizophrenia, bipolar disorder, and autism. For the cognitive tests, only polygenic risk for autism was related with higher cognitive ability, in agreement with a previous study.^25^ Genes related to bipolar disorder were negatively related to all of the cognitive tests, and genes related to schizophrenia even more so. Previous epidemiological studies indicate that both very high and very low educational achievement is associated with an increased risk of bipolar disorder^26^ and high polygenic risk of schizophrenia and bipolar disorder were recently associated with higher levels of creativity.^27^ The discrepancy between the cognitive and educational results may be explained by the age of the participants: if schizophrenia genes are detrimental to cognitive functioning only later in life, they may have differential effects on educational attainment, which tends to peak before age 30, and the cognitive tests, which were taken in UK Biobank at an average age of 56.9 years. It should also be noted the UK Biobank sample consists of individuals who are on average older than those in most schizophrenia genetic studies, have a higher level of education and are of a higher social class. It is also likely to have been the case that individuals in middle age with a history of serious mental illness will have been less likely than those without such a history to volunteer as UK Biobank participants. These demographic and clinical factors might have contributed to apparently contradictory findings with respect to cognitive function and previous educational attainment.

The Educational Attainment variable demonstrated pleiotropy with 20 of the 24 health-related variables, indicating that the genetic variants that collectively act to facilitate an individual’s progress through the educational system to degree level make important contributions to many important health outcomes. One explanation for this is that the educational attainment variable shows the greatest degree of pleiotropy with general cognitive ability in childhood with a genetic correlation of 0.906. This could indicate that it is genes related to cognitive ability early in life that are responsible for the pleiotropy with health variables: educational attainment, therefore, might act as a proxy phenotype for general cognitive ability, as others have demonstrated.^8^

The significant genetic correlations across traits enabled the use of polygenic profile scores to predict phenotypic cognitive variance in the UK Biobank sample. The amount of variance explained by the polygenic risk profiles for each UK Biobank cognitive phenotype is small, as would be expected by the fact that not all SNPs were genotyped, and those that were do not necessarily accurately tag the causal genetic variants. The multivariate polygenic risk score analyses showed that additional variance can be accounted for when the polygenic liabilities of multiple disorders and traits are combined; this implies that there are risk alleles unique to each disorder and trait that affect cognitive and educational traits.

The results of the present study are supported by a previous study examining the genetic associations between polygenic profile scores for psychiatric and cognitive traits, and many phenotypic traits, showing comparable directions of effect^28^. However, due to the much smaller sample size (3000 versus 112 000 in the present study), this study yields insufficient power to detect several of the associations found in the present study. That same study also supported the results of the LD regression analyses of the present study between several psychiatric and cognitive traits, but the following traits show novel associations with cognitive ability and educational attainment in the present study: coronary artery disease, ischaemic stroke, infant head circumference and years of education.

The polygenic profile analysis replicated previous, smaller studies that showed associations between higher cognitive function and higher polygenic risk for autism^25^ and lower polygenic risk for schizophrenia^18^ and stroke^19^. We did not replicate the previous finding that higher cognitive function is associated with higher Type 2 diabetes genetic risk^29^, but we did find that higher Type 2 diabetes genetic risk is associated with decreased likelihood of obtaining a college degree. Unlike a previous small, underpowered, study^20^ we found that higher polygenic risk for Alzheimer’s disease is associated with lower cognitive function.

To the extent that these genetic associations between cognitive and health measures are explained by shared genetic influences, they support the theoretical construct of bodily system integrity^30^. System integrity was formulated as a latent trait which is manifest as individual differences in how effectively people meet cognitive and health challenges from the environment, and which has some genetic aetiology. Whereas it is recognised that some illnesses will cause changes in cognitive functions, system integrity suggests in addition that there is shared variance in how well different complex bodily systems operate and that this underlies correlations between higher cognitive functioning and good health and longevity.

The present study has a number of strengths. First, the large sample size of UK Biobank (*N* > 100,000) affords powerful, robust tests of genetic association. Second, the participants took identical cognitive tests, which were always administered in the same computerised fashion, reducing any potential bias due to heterogeneity in test content and administration. Third, all of the UK Biobank genetic data were processed in a consistent matter, on the same platform and at the same location. Fourth, our use of summary data from a large number of international GWAS studies allowed a comprehensive and detailed examination of shared genetic aetiology with cognitive ability across a wide range of health-related phenotypes, producing many of the first estimates of the genetic correlation between traits.

The present study has some limitations. The three cognitive tests were brief, bespoke measures. However, they covered three major cognitive domains (reasoning, processing speed, and memory), showed acceptable internal consistency, and had validity in that they showed the expected correlations with one another and with age and educational attainment.^31,32^ The Verbal-numerical reasoning test has types of item that are the same as those found in tests of general cognitive ability. In addition, Verbal-numerical Reasoning and Educational Attainment showed strong genetic correlations (0.812 and 0.906, respectively) with childhood general cognitive function. This suggests that the variance in these traits is largely the product of the same genetic variants that underpin general cognitive function. The GWAS studies we curated to perform LD score regression and extract the polygenic profile scores were often consortia studies, involving meta-analyses across datasets with substantial heterogeneity in sample size, genome-wide imputation quality, and phenotypic measurement. We expect that, with larger and more consistent independent datasets, we would be able to use the polygenic profile scores to predict more variance in cognitive test performance. Some of the GWAS consortia studies did not have ethical approval to be used for genetic correlation and polygenic profile scoring analyses associated with education, meaning that we were unable to estimate a few correlations. Clustering in genetic population structure meant that we restricted the genotyped samples to individuals of white British ancestry. Our results thus need to be replicated in large samples of different genetic backgrounds; the sample sizes available in UK Biobank were not large enough for us to model, with adequate power, data from UK Biobank individuals of other ancestries.

The best method for showing pleiotropy using GWAS data would be to obtain the genome wide significant hits from two GWAS and correlate the effect sizes. However, there are two reasons why this is suboptimal at present. The first reason is that many significant SNP hits are needed in multiple GWAS data sets which is currently not possible. The second reason is that GCTA has shown that many true associations do not reach statistical significance due to low power, therefore SNPs that do not attain statistical significance should also be considered. Both LD regression and polygenic profile score analyses provide the opportunity to use the full GWAS output to examine pleiotropy.

Because the optimum number of SNPs used to generate a particular polygenic profile can differ between traits, we created five profile scores per physical and mental health trait and tested each in a regression against each of the UK Biobank cognitive traits to determine the score that explained the greatest variance in each cognitive trait. We found that these did differ between different trait combinations, suggesting that the amount of shared genetic aetiology differs between different pairs of traits. However, as pleiotropy was quantified using LD score regression to perform a single test for each pair of phenotypes, the multiple testing problems associated with the polygenic profile score method did not confound the estimates of pleiotropy shown here. Whereas the estimate of phenotypic variance explained by the polygenic profile method was small, this should be considered as the minimum estimate of the variance explained. Due to pruning SNPs in LD, the PGRS method makes the assumption of a single causal variant being tagged in each LD block considered. If this assumption is not true for the phenotypes considered, the proportion of variance explained will be underestimated here.

## Conclusion

It is notable that, a short while ago, a single result from the dozens of reported here would have been considered a major novel finding and reported as a study in itself.^17^ With so many findings, it has not been possible fully to discuss their implications. For example, the genetic associations between infant head circumference and intracranial volume with Educational Attainment and verbal-numerical reasoning are important in themselves, as are many other cognitive-mental health and cognitive-physical health associations. Taken all together, these results provide a resource that advances the study of aetiology in cognitive epidemiology substantially.

## Acknowledgments

This research has been conducted using the UK Biobank Resource. The work was undertaken in The University of Edinburgh Centre for Cognitive Ageing and Cognitive Epidemiology, part of the cross council Lifelong Health and Wellbeing Initiative (MR/K026992/1). Funding from the BBSRC and Medical Research Council (MRC) is gratefully acknowledged.

